# Impaired chloroplast positioning affects photosynthetic capacity and regulation of the central carbohydrate metabolism during cold acclimation

**DOI:** 10.1101/2020.05.06.080531

**Authors:** Anastasia Kitashova, Katja Schneider, Lisa Fürtauer, Laura Schröder, Tim Scheibenbogen, Siegfried Fürtauer, Thomas Nägele

## Abstract

Photosynthesis and carbohydrate metabolism of higher plants need to be tightly regulated to prevent tissue damage during environmental changes. The intracellular position of chloroplasts changes due to a changing light regime. Chloroplast avoidance and accumulation response under high and low light, respectively, are well known phenomena, and deficiency of chloroplast movement has been shown to result in photodamage and reduced biomass accumulation. Yet, effects of chloroplast positioning on underlying metabolic regulation are less well understood. Here, we analysed photosynthesis together with metabolites and enzyme activities of the central carbohydrate metabolism during cold acclimation of the *chloroplast unusual positioning 1* (*chup1*) mutant of *Arabidopsis thaliana*. We compared cold acclimation under ambient and low light and found that maximum quantum yield of PSII was significantly lower in *chup1* than in Col-0 under both conditions. Our findings indicated that net CO_2_ assimilation in *chup1* is rather limited by biochemistry than by photochemistry. Further, cold-induced dynamics of sucrose phosphate synthase differed significantly between both genotypes. Together with a reduced rate of sucrose cycling derived from kinetic model simulations our study provides evidence for a central role of chloroplast positioning for photosynthetic and metabolic acclimation to low temperature.

## Introduction

Tight regulation and balance of photosynthetic primary and secondary reactions are a prerequisite for efficient CO_2_ fixation, biomass accumulation and stress tolerance of higher plants. Particularly under changing environmental conditions, stabilization of this balance is essential for prevention of tissue damage due to cytotoxic accumulation of reactive oxygen species (ROS). Intracellular distribution of chloroplasts changes in response to changing light intensity. Chloroplast movement plays an important role in protection of photosynthetic machinery against high light intensities and increases photosynthetic efficiency under low light (Wada, 2013). Typically, high light induces an avoidance response where chloroplasts are located at the side-walls of palisade cells to minimize light absorption. Under low light, chloroplasts re-distribute to the upper and lower side of the cells to maximize light absorption (accumulation response; schematic overview in (Wada, 2013)).

The cytoplasmic protein CHLORPLAST UNUSUAL POSITIONING 1 (CHUP1) localizes to the chloroplast surface, contains an actin binding domain and has been shown to be essential for chloroplast movement (Oikawa *et al.*, 2003, Oikawa *et al.*, 2008). It was demonstrated that CHUP1 interacts with actin, independent of its filamentation status, and further binds to profilin which is an actin modifying protein (Schmidt von Braun and Schleiff, 2008). On gene expression level, no alteration of other components involved in chloroplast positioning was observed in a *CHUP1* deletion mutant (Schmidt von Braun and Schleiff, 2008). From their gene expression data, the authors concluded that the observed genetic effect rather resulted from altered positioning of chloroplasts and peroxisomes, which was shown to be coupled before (Oikawa *et al.*, 2003), than from alteration of genes encoding for components of light perception and chloroplast movement. Under short-day growth conditions, i.e. 8h light phase, *chup1* plants displayed reduced photosynthesis and biomass accumulation over a wide range of light intensities, i.e. 50 – 300 μmol m^−2^ s^−1^ (Gotoh *et al.*, 2018). In their study, Gotoh and co-workers compared different mutants affected in chloroplast positioning and finally concluded that trade-offs of biomass production and photoprotection are essentially driven by chloroplast movement under changing growth light.

With decreasing temperature, changes in light intensity become an increasing challenge for plants. During the multigenic process of cold acclimation, which is induced by exposure of higher plants to low but non-freezing temperature, numerous molecular and physiological processes are adjusted to enhance freezing tolerance and to minimize cellular and tissue damage, e.g. caused by ROS production (Levitt, 1980, Xin and Browse, 2000, Dreyer and Dietz, 2018). Cold acclimation plays an important role in ecology and evolution of many plant species of the temperate zone due to the significant impact of low temperature on their range boundaries (Hoffmann, 2002). It affects gene expression, translational, post-translational and metabolic regulation (Wang *et al.*, 2017, Bahrani *et al.*, 2019, Fürtauer *et al.*, 2019, Liu *et al.*, 2019). Further, composition and activation state of the photosynthetic apparatus is adjusted within a process termed photosynthetic acclimation, in which, however, many of the involved molecular mechanisms still remain elusive (Herrmann *et al.*, 2019). Low temperature is generally accompanied by a reduced membrane fluidity which affects, e.g. the repair cycle of the D1 protein (Aro *et al.*, 1990). Additionally, low temperature significantly affects enzyme activities of the Calvin-Benson cycle which needs to be counteracted during cold stress response and acclimation to prevent imbalance between primary and secondary photosynthetic reactions (Khanal *et al.*, 2017). Absorption of excessive sunlight by the light-harvesting complex leads to ROS generation which damages the photosystems, particularly PSII, and finally leads to photoinhibition (Gururani *et al.*, 2015). Yet, while particularly under low temperature excessive light is harmful for plants, a combination of low temperature and light is essential for enhancing freezing tolerance in *Arabidopsis thaliana* (Wanner and Junttila, 1999).

Comparing temperature effects on photosynthesis in natural Arabidopsis accessions previously revealed that limitation of photosynthesis by triose-phosphate utilization was less under low temperature than under high temperature comparing low vs. high irradiance (Pons, 2012). Together with many other studies, this observation clearly emphasizes the essential role of tight regulation of photosynthesis and carbohydrate metabolism. Sucrose (Suc) metabolism is well known to play a central role in photosynthetic acclimation (Strand *et al.*, 2003, Stitt *et al.*, 2010, Nägele *et al.*, 2012, Herrmann *et al.*, 2019). In the light, photosynthetic CO_2_ fixation results in triose phosphates which are substrate for starch biosynthesis in the chloroplast and Suc biosynthesis in the cytosol. Key regulators of Suc biosynthesis are cytosolic fructose-1,6-bisphosphatase (cFBPase) and sucrose phosphate synthase (SPS) (Strand *et al.*, 2000). Suc is the central transport sugar in many plant species, represents an osmotically active solute and is involved in sugar signalling (Ruan, 2014). Supportingly, recent findings suggest that Suc export from the chloroplast into the cytosol is involved in cold acclimation and is required for development of maximal freezing tolerance (Patzke *et al.*, 2019). Also, in our own studies we demonstrated that Suc compartmentation and invertase-driven cleavage in cytosol and vacuole significantly affect cold stress response and stabilization of photosynthesis (Nägele and Heyer, 2013, Weiszmann *et al.*, 2018).

In the present study we aimed to reveal the effect of defective chloroplast positioning on photosynthesis and regulation of carbohydrate metabolism during cold acclimation under ambient and low light in *Arabidopsis thaliana*.

## Materials and Methods

### Plant material and growth conditions

Plants of *Arabidopsis thaliana* accession Col-0 and the homozygous *chup1* T-DNA insertion line SALK_129128C (locus At3g25690; validated by PCR) were grown on a 1:1 mixture of GS90 soil and vermiculite in a climate chamber under short-day conditions (8h/16h light/dark; 100 μmol m^−2^sec^−1^; 22°C/16°C; 60% relative humidity). After 6 weeks, plants were transferred to the greenhouse and grown under long day conditions (16h/8h light/dark). After 5 days in the greenhouse, plants were either (i) sampled at midday, i.e. after 8h in the light, or (ii) transferred to a cold room for cold acclimation at 4°C under long day conditions (16h/8h light/dark), with either PAR 100 μmol m^−2^ sec^−1^ (samples ambient light/low temperature, AL/LT) or PAR 20 μmol m^−2^ sec^−1^ (samples low light/low temperature, LL/LT). After 7 days at 4°C, plants were sampled at midday after 8h in the light. One sample consisted of 3 leaf rosettes which were immediately frozen in liquid nitrogen and stored at −80°C until further use. For photosynthesis measurements, plants were grown with the same protocol.

### Net photosynthesis and chlorophyll fluorescence measurements

Maximum quantum yield of PSII (Fv/Fm) was determined after 15 min of dark adaptation by supplying a saturating light pulse. Dynamics of quantum efficiency of PSII (ΦPSII), electron transport rate (ETR), photochemical (qP), non-photochemical quenching (qN) and rates of net CO_2_ assimilation, i.e. net photosynthesis (NPS), were determined within light response curves by stepwise increase of photosynthetically active radiation (PAR) from 0 to 1200 μmol m^−2^ s^−1^ in 5 min intervals. Additionally, all parameters were recorded within CO_2_ response curves where CO_2_ concentration was stepwise increased from 50ppm to 1200ppm under constant PAR intensity of 1200 μmol m^−2^ s^−1^. All measurements were performed using a WALZ® GFS-3000FL system equipped with measurement head 3010-S and Arabidopsis chamber 3010-A (Heinz Walz GmbH, Effeltrich, Germany, https://www.walz.com/). NPS and chlorophyll fluorescence parameters of control plants were determined at ambient temperature (22°C), parameters of AL/LT and LL/LT plants were determined at 4°C.

### Quantification of amount of starch, soluble carbohydrates and hexose phosphates

Starch amount was determined as described previously with slight modification (Nägele *et al.*, 2012). 400μl of 80% EtOH were added to ground plant material and incubated at 80°C for 30min. After centrifugation, the supernatant was transferred into a new tube and extraction was repeated. Supernatants (80% EtOH) were pooled, dried in a desiccator and used for quantification of soluble carbohydrates. Starch containing pellets were incubated with 500μl 0.5N NaOH at 95°C for 60min before slight acidification with 500μl 1M CH_3_COOH. The starch hydrolysate was digested with amyloglucosidase for 2h at 55°C. Glucose content, derived from starch digestion, was photometrically determined by a glucose oxidase/peroxidase/o-dianisidine reaction at 540nm.

Soluble sugars, i.e. glucose, fructose and sucrose, were determined from dried EtOH extracts. Sucrose was quantified using an anthrone assay after incubation with 30% KOH at 95°C. For the assay, anthrone was dissolved in 14.6M H_2_SO_4_ (0.14% w/v), incubated with the samples for 30min at 40°C before absorbance was determined at 620nm.

Glucose amount was determined using a coupled hexokinase/glucose 6-phosphate dehydrogenase assay resulting in NADPH+H^+^ which was determined photometrically at 340nm. For fructose quantification, phosphoglucoisomerase was added to the reaction mixture after glucose determination.

Glucose 6-phosphate (G6P) and fructose 6-phosphate (F6P) were extracted and photometrically quantified using enzyme cycling-based assays as described earlier (Gibon *et al.*, 2002). In brief, hexose phosphates were extracted using trichloroacetic acid (TCA) in diethyl ether (16% w/v), washed with 16% (w/v) TCA in 5mM EGTA and neutralised with 5M KOH/1M triethanolamine. Enzymatic reactions catalysed the equimolar interconversion of G6P and F6P into NADPH+H^+^ which was finally detected photometrically from slopes of a cyclic reaction yielding a formazan dye from thiazolyl blue (MTT) at 570nm.

### Quantification of enzyme activities

Activity of all enzymes was determined under substrate saturation (v_max_).

Activity of sucrose phosphate synthase (SPS) was determined using the anthrone assay as described earlier (Nägele *et al.*, 2012). Frozen leaf tissue was suspended in extraction buffer containing 50mM HEPES-KOH (pH 7.5), 10mM MgCl_2_, 1mM EDTA, 2.5mM DTT, 10% (v/v) glycerine and 0.1% (v/v) Triton-X-100. After incubation on ice, extracts were incubated for 30min at 25°C with reaction buffer containing 50mM HEPES-KOH (pH 7.5), 15mM MgCl_2_, 2.5mM DTT, 35mM UDP-glucose, 35mM F6P and 140mM G6P. Reactions were stopped by adding 30%KOH and heating the samples to 95°C. Sucrose was determined photometrically after incubation with anthrone in H_2_SO_4_ as described above and blanks (stopped before reaction) were subtracted.

Gluco- and fructokinase activities were determined photometrically at 340nm recording the slopes of NADP^+^ reduction to NADPH+H^+^ (Wiese *et al.*, 1999). In brief, frozen leaf tissue was suspended in extraction buffer containing 50mM Tris-HCl (pH 8.0), 0.5mM MgCl_2_, 1mM EDTA, 1mM DTT and 1% (v/v) Triton-X-100. Following incubation on ice, reaction was started with a reaction buffer containing 100mM HEPES-KOH (pH 7.5), 10mM MgCl_2_, 2mM ATP, 1mM NADP, 0.5U G6PDH and either 5mM glucose for glucokinase measurement or 5mM fructose and 0.5U PGI for fructokinase measurement.

Neutral (nInv), acidic (aInv) and cell wall-bound (cwInv) invertase activities were determined as described before with slight modifications (Nägele *et al.*, 2010). Frozen leaf tissue was homogenized in extraction buffer containing 50mM HEPES-KOH (pH 7.5), 5mM MgCl_2_, 2mM EDTA, 1mM phenylmethylsulfonylfluoride (PMSF), 1mM DTT, 10% (v/v) glycerine and 0.1% (v/v) Triton-X-100. After incubation on ice, samples were centrifuged at 4°C and supernatants were transferred to a new tube. Pellets contained cell wall-bound invertase and were re-suspended in extraction buffer. Soluble, i.e. nInv and aInv, invertase activities were determined from supernatants, cwInv activity was determined from resuspended pellets. Activity of nInv was determined using a reaction buffer with pH 7.5 (20mM HEPES-KOH, 100mM sucrose), aInv and cwInv activities were determined at pH 4.7 (20mM sodium acetate, 100mM sucrose). After incubation of extract with reaction buffers reactions were stopped at 95°C and glucose built from invertase reactions was photometrically determined by a glucose oxidase/peroxidase/o-dianisidine assay at 540nm.

Maximal activity of glucose 6-phosphate dehydrogenase (G6PDH) was determined as reported earlier (Fahrendorf, Ni et al. 1995) with slight modification. Ground plant material was suspended in extraction buffer consisting of 0.05 M Tris HCl (pH 8.0), 0.3 M NaCl, 0.1 mM benzamidine, 0.1 mM PMSF, 1 mM DTT and 0.1% (v/v) Triton-X-100. After incubation on ice and centrifugation at 4°C, supernatants were incubated with assay buffer containing 0.1 M Tris HCl (pH 8.0) and 0.4 mM NADP+. After pre-incubation at 30°C for 10 min, G6P was added and reduction of NADP+ by G6PDH was measured photometrically at 340 nm at 30 °C for 10 min.

Phosphoglucose isomerase (PGI) extraction and maximal activity quantification was based on previously described methods with slight modification (Jones, Pichersky et al. 1986, Kunz, Zamani-Nour et al. 2014). Ground plant material was suspended in extraction buffer containing 0.05 M Tris HCl (pH 8.0), 0.5 mM MgCl_2_, 1 mM EDTA, 2.5mM DTT and 1% (v/v) Triton-X-100. The suspension was incubated on ice, followed by centrifugation at 4°C. PGI activity was estimated from supernatants in a coupled reaction with G6PDH by photometrically measuring reduction of NADP+ at 340 nm. For this, supernatants were incubated with assay buffer consisting of 100 mM HEPES-KOH (pH 7.5) and 10mM MgCl_2_. After addition of 2μl of 100mM NADP^+^ and 1 U G6PDH, the solution was pre-incubated at 30°C for 10 min. The reaction was initiated by addition of 20 μl of 50 mM F6P (or 20 μl of ddH_2_O for blanks), followed by recording absorbance at 340 nm at 30°C for 10 min. The linear range of recorded kinetic was interpolated and the slope was used to calculate maximal activity of PGI.

### Sample preparation and sectioning for microscopy analysis

Primary leaves of *Arabidopsis thaliana*, Col-0 and *chup1*, were cut into 1 mm square pieces and fixed in fixation buffer (75 mM sodium cacodylate, 2 mM MgCl_2_, pH 7) substituted with 2.5% (v/v) glutaraldehyde for 2 days at 4°C. The leave pieces were washed three times at room temperature using fixation buffer and post-fixed with 1% (w/v) osmium tetroxide for 2 hours. Subsequent to washing with fixation buffer and water, the samples were stained *en* bloc with 1% (w/v) uranyl acetate in 20% (v/v) acetone for 30 minutes. Samples were dehydrated in a series of graded acetone and embedded in Spurr’s resin. Samples were either ultra-thin sectioned (thickness, 60 nm) for TEM (transmission electron microscope) analysis or semi-thin sectioned (thickness, 2 μm) for LM (light microscope) analysis. For sectioning a diamond knife was used on a Reichert Ultracut-E ultramicrotome. Ultra-thin sections were mounted on collodium-coated copper grids, post-stained with lead citrate (80 mM, pH 13) and examined with an EM 912 transmission electron microscope (Zeiss, Oberkochen, Germany) equipped with an integrated OMEGA energy filter operated in the zero-loss mode at 80 kV. Images were acquired using a 2k x 2k slow-scan CCD camera (Tröndle Restlichtverstärkersysteme, Moorenweis, Germany). Semi-thin sections mounted on glass slides and examined using a Zeiss Axiophot light microscope. Images were acquired using a SPOT insight 2 MP CCD color digital camera.

### Statistics and mathematical modelling

For statistical data evaluation we used the free software environment R Version 3.6.1 (https://www.r-project.org/) (R Core Team, 2019) and RStudio Version 1.2.5019 (https://rstudio.com/) (RStudio Team, 2019). Statistics comprised analysis of variance (ANOVA) and Tukey’s honestly significant difference post-hoc test. Mathematical modelling was performed in MATLAB® R2019b (www.mathworks.com) with the IQM Tools developed by IntiQuan (https://www.intiquan.com/). Model structure (ODEs), enzyme kinetic equations and kinetic parameter values are provided in the supplements (Supplementary Table IV – VI). Rates of net photosynthesis (‘rNPS’) were derived from interpolation of photosynthesis measurements, i.e. net CO_2_ assimilation rates (interpolation model: f(x) = a*exp(b*x) + c*exp(d*x)). For simulation of sucrose cleavage rates, experimentally quantified invertase activities of nInv, aInv and cwInv were summed up (model parameter ‘vmax_inv’). Rates of net starch synthesis (‘rStarch’) were estimated from the amount measured after 8h in the light (i.e. amount/8h). For simulation at low temperature (AL/LT and LL/LT), quantified vmax values were temperature corrected to 4°C applying the van’t Hoff rule (Eq. 1) with a Q_10_ factor of 2.5 (Reyes *et al.*, 2008).

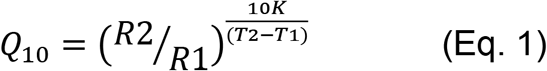

Ranges of kinetic parameters which were not experimentally quantified in the present study, i.e. K_M_ and K_i_ values, were derived from our previous studies and further literature (Schnarrenberger and Oeser, 1974, Scharte *et al.*, 2009, Nägele *et al.*, 2010, Nägele *et al.*, 2012, Fürtauer and Nägele, 2016, Weiszmann *et al.*, 2018).

## Results

### The CHUP1 mutation significantly affects maximum quantum yield of PSII under low temperature and rates of net CO_2_ fixation under high light

Affected chloroplast positioning in *chup1* (see Supplementary Figures S1 and S2) had significant effects on photosynthesis both under ambient and low temperature. While Fv/Fm did not differ between *chup1* and Col-0 under control conditions, it was lower in *chup1* than in Col-0 at 4°C under both ambient and low light (Fig. 1; AL/LT: p=0.07; LL/LT: p<0.05, ANOVA). Interestingly, in both genotypes Fv/Fm was significantly higher under LL/LT (>0.82) than under control (~0.815) conditions (p < 0.001, ANOVA). In summary, maximum quantum yield of PSII did not differ between both genotypes under control conditions but was lower in *chup1* than in Col-0 after 7 days at 4°C under both low (20 μmol m^−2^ s^−1^) and ambient light (100 μmol m^−2^ s^−1^).

**Figure 1.**
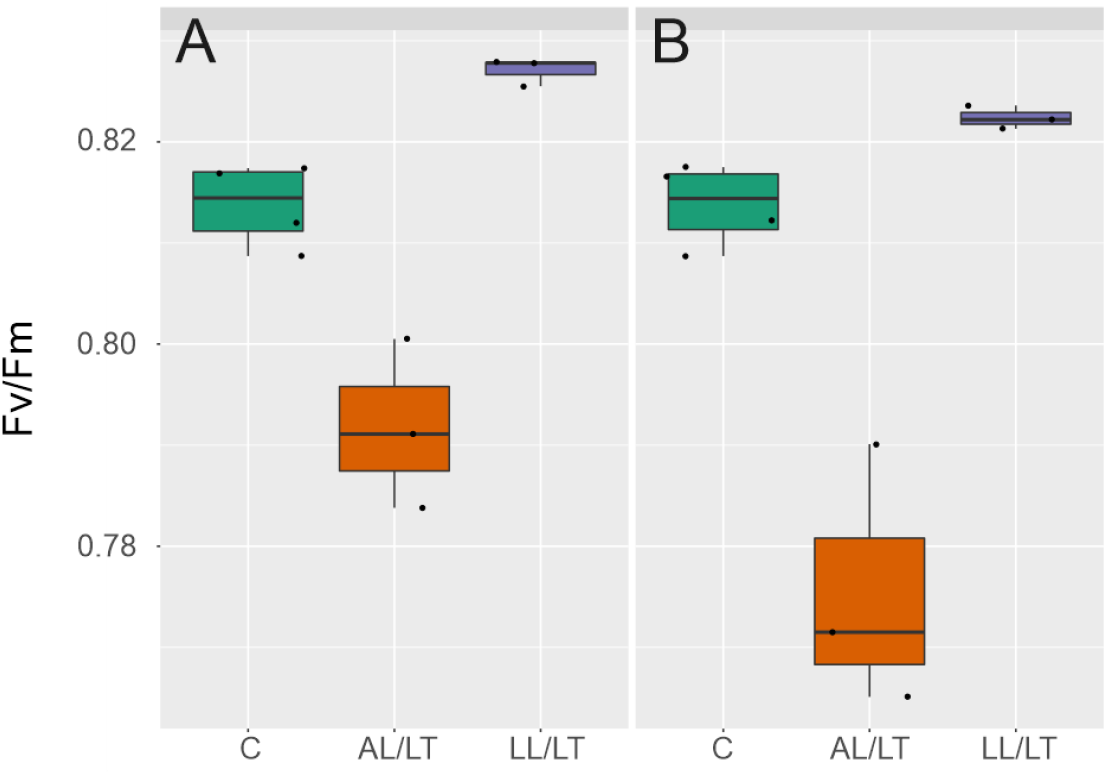
Maximum quantum yield of PSII in Col-0 and *chup1*. (A) Fv/Fm in Col-0 and (B) in *chup1* under control (green box, C), ambient light/low temperature (orange box, AL/LT, 100 μmol m^−2^ s^−2^) and low light/low temperature (purple box, LL/LT, 20 μmol m^−2^ s^−2^). In addition to coloured box-and-whisker plots single measurements are indicated by black dots.

Rates of net CO_2_ assimilation were recorded together with chlorophyll fluorescence parameters within light response curves and CO_2_ response curves (Fig. 2, Supplementary Figures S3 and S4). Analysis of variance (ANOVA) revealed a highly significant condition effect (p < 0.001) on net CO_2_ fixation rates of both genotypes separating control vs. AL/LT and control vs. LL/LT across a wide range of PAR intensity and CO_2_ concentration (Fig. 2 A-D). No significance was observed between AL/LT and LL/LT neither in light response curves nor in CO_2_ response curves. Further, under high PAR intensity, i.e. 1200 μmol m^−2^ s^−1^, and at ambient temperature (22°C) *chup1* had a significantly lower net CO_2_ fixation rate than Col-0 (p<0.001, ANOVA; a summary of all ANOVA results in provided in the supplements, Supplementary Tables I–III). Local polynomial regression under control conditions (green lines in Fig. 2) indicated that net CO_2_ assimilation rates of *chup1* were near to saturation at 1200 μmol m^−2^ s^−1^ (Fig. 2 B) which was not observed for Col-0 (Fig. 2 A). Under high CO_2_ concentration, i.e. 900 and 1200ppm, and high irradiance (1200 μmol m^−2^ s^−1^), net CO_2_ assimilation rates did not significantly differ between Col-0 and *chup1* (Fig. 2 C, D). Instead, Col-0 had significantly higher CO_2_ fixation rates than *chup1* at 200ppm and 400ppm (ANOVA, p<0.01). After 7d at 4°C, CO_2_ assimilation rates were found to neither differ between genotypes nor between light conditions within a genotype, i.e. AL/LT and LL/LT.

**Figure 2.**
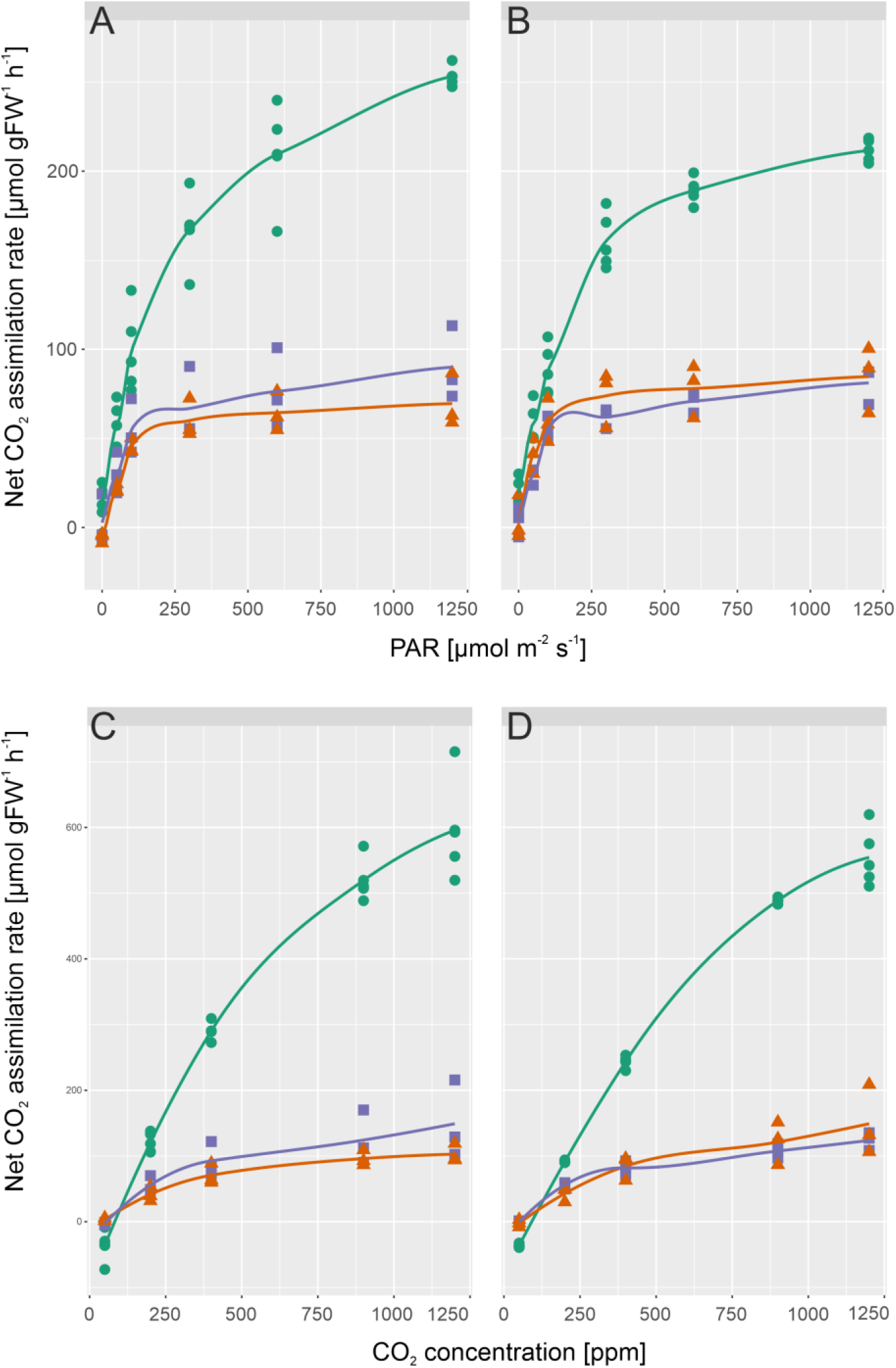
Rates of net CO_2_ assimilation in Col-0 and *chup1* as a function of irradiance and CO_2_ concentration. **(A)** Rates of net CO_2_ assimilation in Col-0 under control (green circles), AL/LT (orange triangles) and LL/LT (purple squares) as a function of PAR, **(B)** Rates of net CO_2_ assimilation in *chup1* under control (green circles), AL/LT (orange triangles) and LL/LT (purple squares) as a function of PAR, **(C)** Rates of net CO_2_ assimilation in Col-0 under control (green circles), AL/LT (orange triangles) and LL/LT (purple squares) as a function of CO_2_ concentration, **(D)** Rates of net CO_2_ assimilation in *chup1* under control (green circles), AL/LT (orange triangles) and LL/LT (purple squares) as a function of CO_2_ concentration. Symbols represent measurements of independent biological replicates (n≥3). Lines represent a local polynomial regression. Measurements of control samples were performed at 22°C, measurements of AL/LT and LL/LT at 4°C.

Light and CO_2_ response curves of chlorophyll fluorescence parameters were recorded together with net CO_2_ assimilation rates (Supplementary Figures S3 and S4) revealing no significant difference between Col-0 and *chup1* (Supplementary Table 1). Yet, only in Col-0 qN was significantly higher under control than under low temperature at high PAR intensity, i.e. 1200 μmol m^−2^ s^−1^ (ANOVA, p<0.05; Supplementary Figure 3 G).

### Low temperature induced accumulation of hexose phosphates is significantly affected in *chup1*

Quantification of central carbohydrates revealed similar amounts of starch and sucrose in Col-0 and *chup1* under all tested conditions (Fig. 3). In both genotypes, starch increased significantly ~10-fold under AL/LT (ANOVA, p< 0.001) and decreased significantly to a level ~0.5 μmol gFW^−1^ under LL/LT (ANOVA, p<0.05; Figure 3 A, B). Similarly, sucrose increased significantly ~2.5-fold under AL/LT (ANOVA, p<0.001) while no significant change was observed under LL/LT neither for Col-0 nor for *chup1* (Fig. 3C, D). Although it is difficult to estimate biological variance from 5 biological replicates, our data indicated a slightly higher variance of starch and sucrose amount in *chup1* than in Col-0, particularly under control and AL/LT.

**Figure 3.**
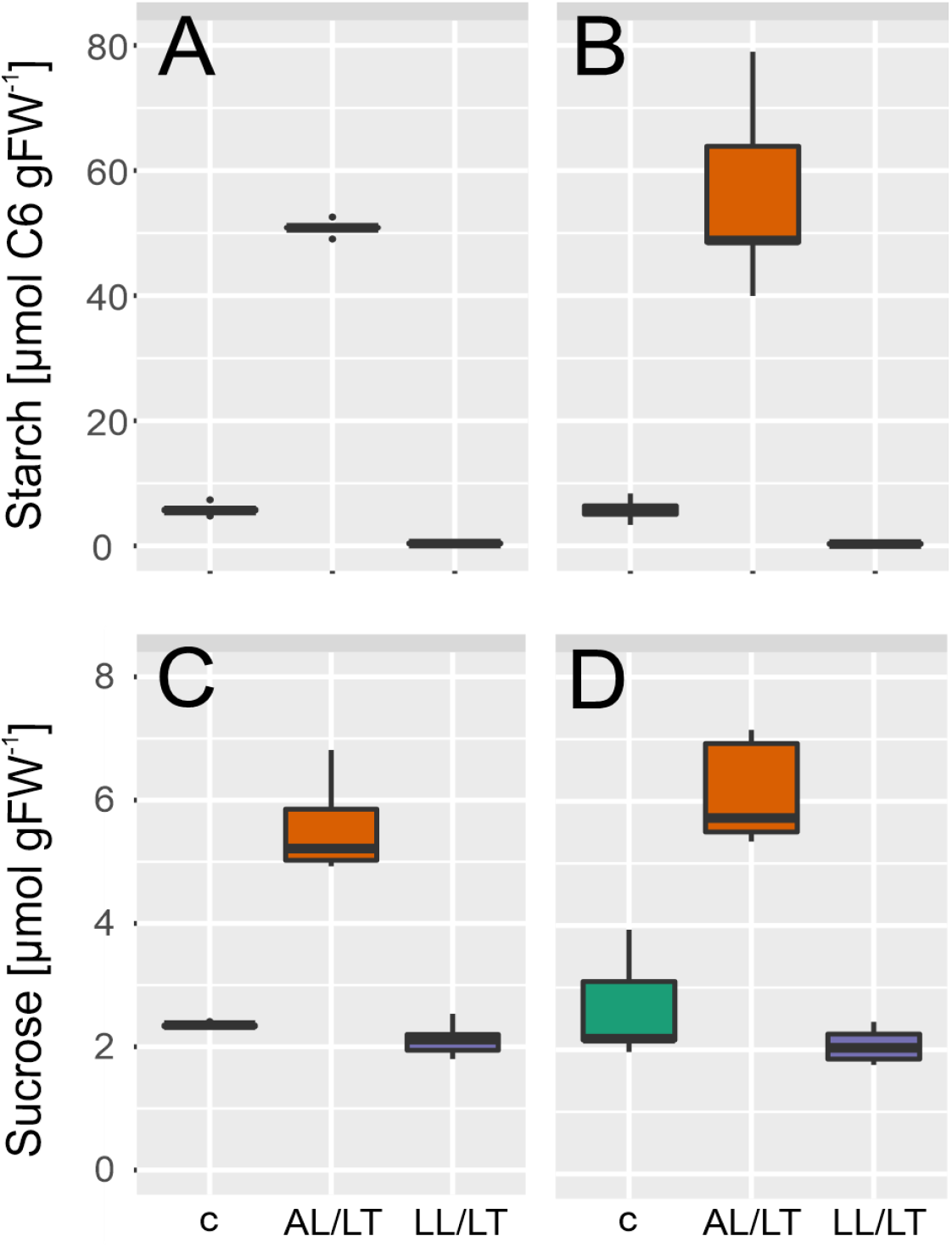
Cold-induced dynamics of starch and sucrose. (**A**) Starch amount in Col-0, (**B**) Starch amount in *chup1*, (**C**) Sucrose amount in Col-0 and (**D**) Sucrose amount in *chup1*. In each panel, left/green: control; middle/orange: AL/LT; right/purple: LL/LT. (n = 5)

Next, we quantified dynamics of free hexoses, i.e. glucose and fructose, and of their phosphorylation products, G6P and F6P (Fig. 4). In both genotypes, glucose and fructose significantly increased under AL/LT (p<0.001) and significantly dropped under LL/LT (p<0.05) compared to the amount under control conditions (Fig. 4 A-D). Neither amount of glucose nor fructose significantly differed between Col-0 and *chup1* under any tested condition. In contrast, dynamics of G6P and F6P revealed significant genotype-effects under low temperature (Fig. 4, E-H). While G6P significantly increased ~3-fold in both genotypes under AL/LT, only in Col-0 we observed a significant difference between G6P amount under AL/LT and LL/LT (Fig. 4 E, F). In *chup1*, G6P amount was similar in LL/LT and AL/LT (Fig. 4 F). Also, dynamics of F6P amount significantly differed between Col-0 and *chup1* where already under control conditions a reduced amount was observed in *chup1* (p=0.06, ANOVA). Under AL/LT, amount of F6P significantly increased ~3-fold in *chup1* while no change was observed in Col-0 (Fig. 4 G, H). Only under LL/LT, F6P amount increased ~1.5-fold in Col-0 (p<0.001). In summary, sucrose/starch ratios and free hexose amounts were similar in both genotypes while hexose phosphates significantly differed in their cold-induced dynamics between Col-0 and *chup1*.

**Figure 4.**
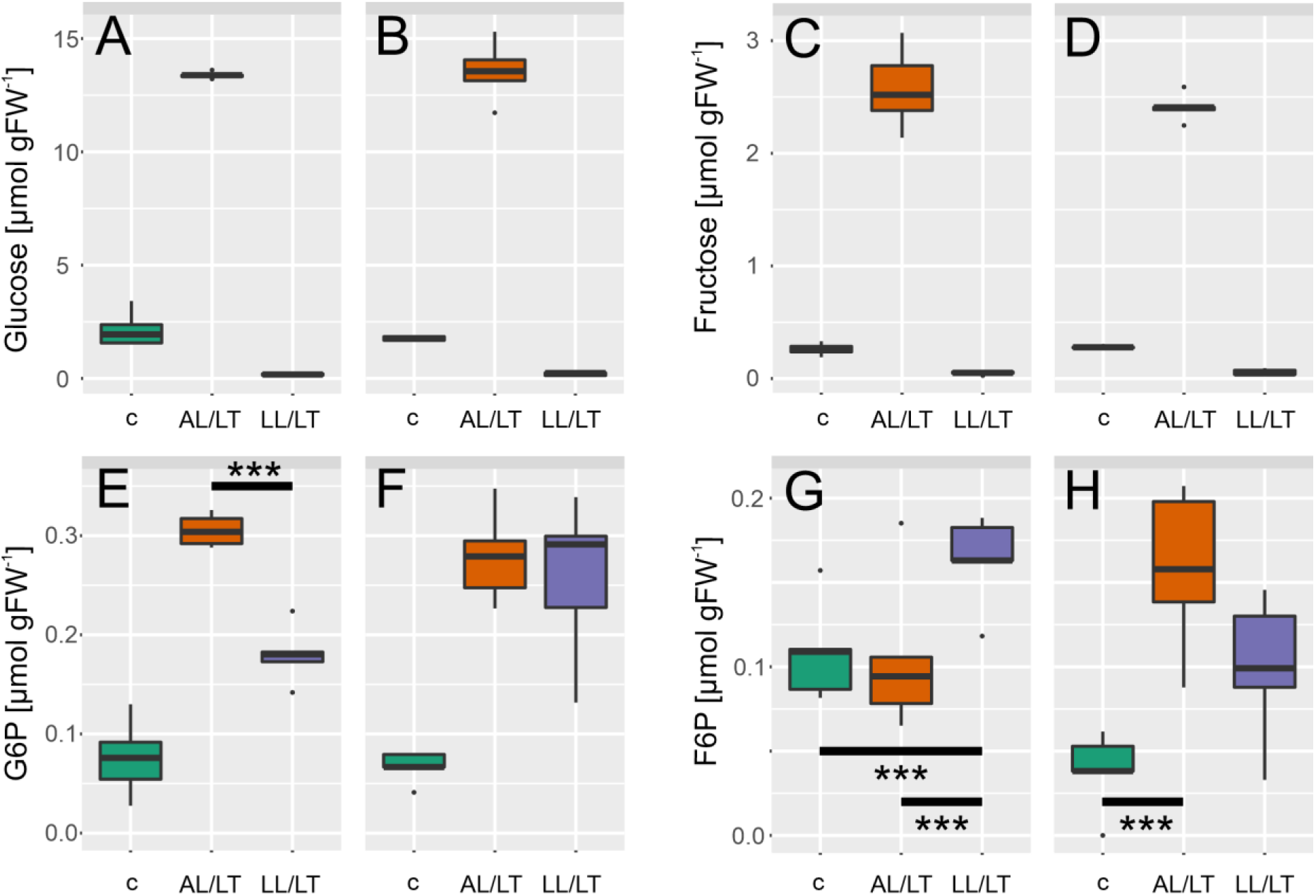
Cold-induced dynamics of hexoses and hexose phosphates. (**A, C, E, G**) amount of substance in Col-0, (**B, D, F, H**) amount of substance in *chup1*. Boxes in each panel, left/green: control; middle/orange: AL/LT; right/purple: LL/LT. Significances are indicated by asterisks but only shown where different between Col-0 and *chup1* (ANOVA, *** p<0.001). A complete overview of significances is provided in the supplements (Supplementary Table I). n = 5.

### Cold-induced dynamics of gluco- and fructokinase activities are significantly reduced in *chup1*

Enzyme activities of the central carbohydrate metabolism were quantified to reveal a potential effect of the CHUP1 mutation on metabolic regulation (Fig. 5). Dynamics of sucrose phosphate synthase (SPS) activity under AL/LT was reduced in *chup1* (Fig. 5 A, B). Most significant effects were observed for gluco-(GlcK) and fructokinase (FrcK) activities under low temperature (Fig. 5 C-F). In Col-0, GlcK activity significantly increased ~3-fold under AL/LT compared to control conditions while no effect was observed in *chup1* (Fig. 5 C, D). Although not significantly, also FrcK increased in Col-0 under AL/LT and this effect was less pronounced in *chup1* (Fig. 5 E, F). Under LL/LT, FrcK activity dropped in both genotypes but only in Col-0 this effect was significant (Fig. 5 E). Activities of neutral, acidic and cell wall-associated invertases were reduced significantly under low temperature in both genotypes (see ANOVA results in Supplementary Table III) and this reduction was most pronounced under AL/LT (Fig.5 G-L). Activities of phosphoglucoisomerase (PGI) and glucose 6-phosphate dehydrogenase revealed similar cold-induced changes in both genotypes with an activity increase under AL/LT and no change or only a slight decrease under LL/LT (Supplementary Figure S5).

**Figure 5.**
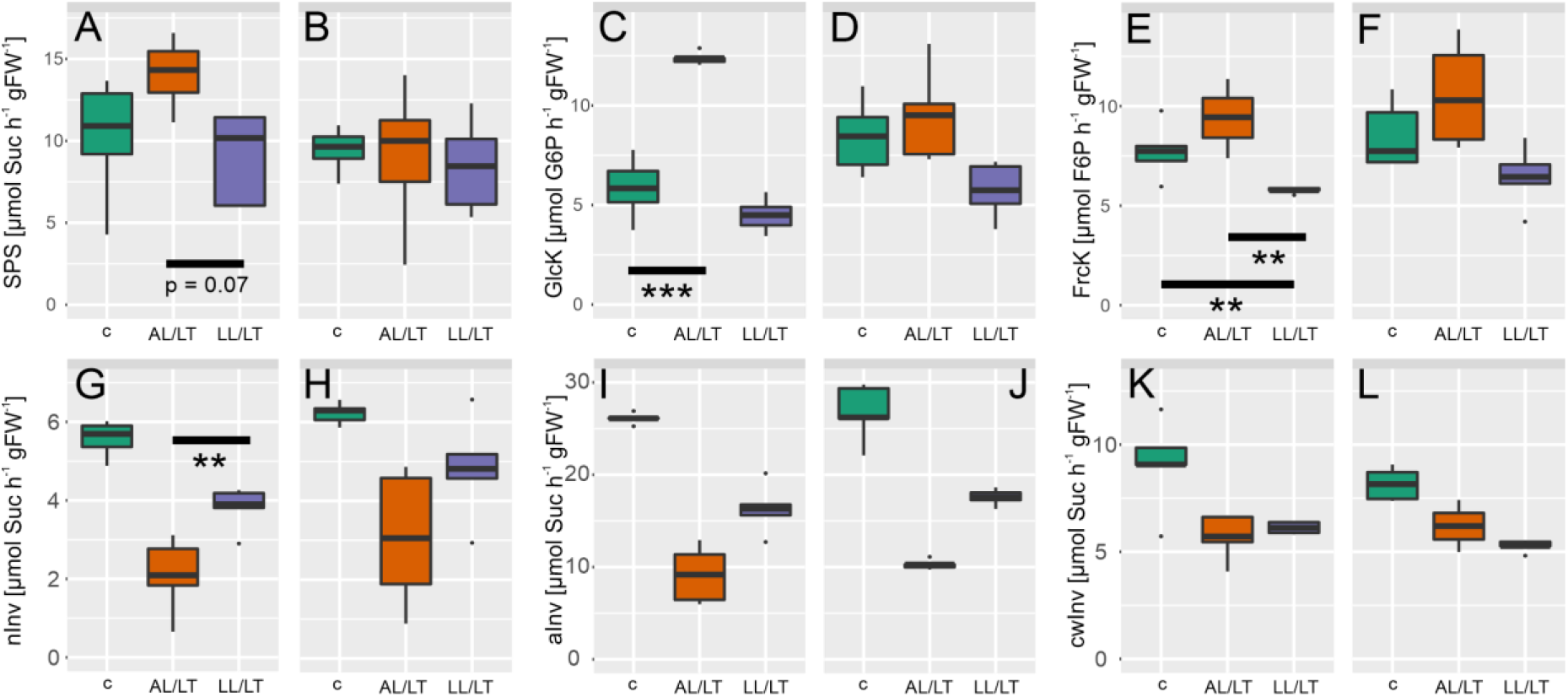
Dynamics of enzyme activities of the central carbohydrate metabolism. **(A, C, E, G, I, K)** enzyme activities in Col-0, **(B, D, F, H, J, L)** enzyme activities in *chup1*. Boxes in each panel, left/green: control; middle/orange: AL/LT; right/purple: LL/LT. SPS: sucrose phosphate synthase; GlcK: glucokinase; FrcK: fructokinase; nInv: neutral invertase; aInv: acidic invertase; cwInv: cell wall-associated invertase. Significances are indicated by asterisks but only shown where different between Col-0 and *chup1* (ANOVA; ** p <0.01; *** p<0.001). A complete overview of significances is provided in the supplements (Supplementary Table I). n = 5.

### Kinetic modelling reveals a shift of reaction rates in *chup1* from hexose phosphorylation into G6P oxidation under low temperature

Metabolite concentrations, net rates of CO_2_ assimilation and enzyme activities were integrated into a mathematical kinetic model. Graphical model structure and identified steady state models for Col-0 and *chup1* under all tested conditions are provided in the supplement (Supplemental Figure S6 and Supplemental Tables IV – VI). For kinetic simulation, a steady state was identified by parameter optimization using temperature-corrected enzyme activities and mean values of quantified metabolites. Particularly under AL/LT, simulations revealed that reactions rates of PGI and G6PDH in Col-0 decreased stronger than in *chup1* (Fig. 6). *Vice versa*, reaction rates of hexose phosphorylation (‘rGLCK’ and ‘rFRCK’) together with sucrose cleavage (‘rINV’) were lower in *chup1* than in Col-0 at AL/LT. Simulated rates of sucrose biosynthesis (‘rSPS’) decreased to similar values in both genotypes. In summary, kinetic simulations suggest that particularly under AL/LT a higher rate of glucose oxidation, catalysed by G6PDH, in *chup1* leads to reduced rates of hexose phosphorylation when compared to Col-0.

**Figure 6.**
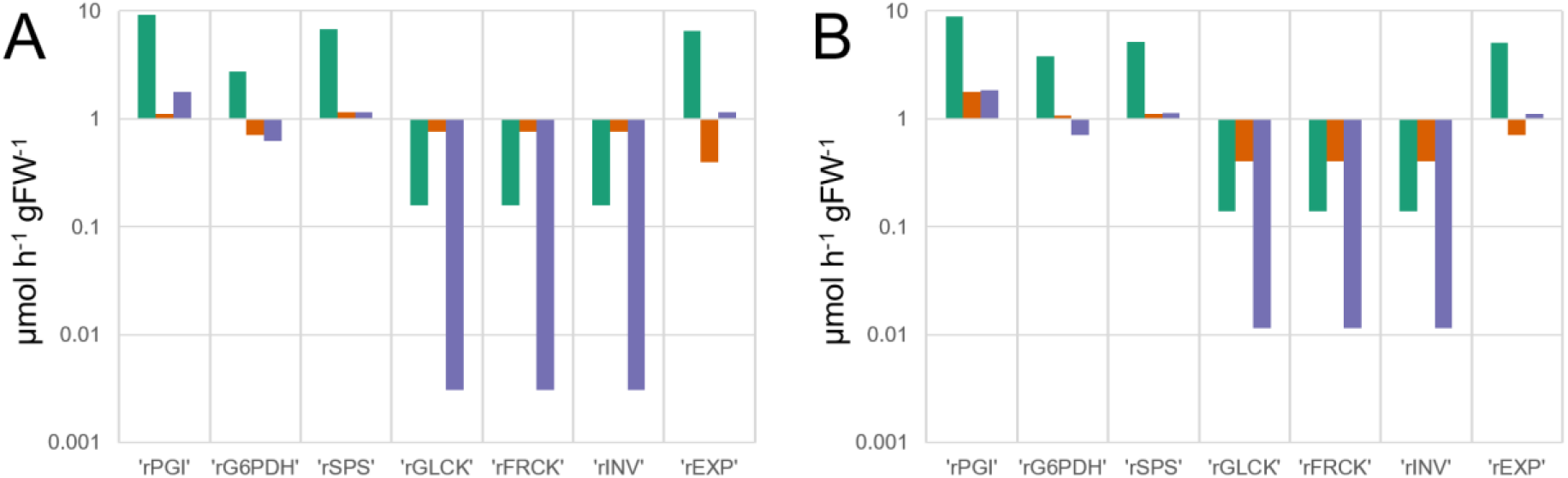
Simulated reaction rates before and after cold acclimation in Col-0 and *chup1*. **(A)** Reaction rates of Col-0 under control, AL/LT and LL/LT. **(B)** Reaction rates of *chup1* under control, AL/LT and LL/LT. Green bars: control; orange bars: AL/LT; purple bars: LL/LT. Simulations were derived from a kinetic model which is explained in the supplements (Supplementary Figure S6, Supplementary Table IV). Enzyme activities were temperature corrected to 4°C for AL/LT and LL/LT simulations.

## Discussion

Chloroplast avoidance movement is a protective mechanism by which plants mitigate photodamage. Under high light intensities chloroplasts typically move to the side walls of cells to minimize the area of potential light absorption. Mutants like *chup1* which are defective in this avoidance movement have been shown to be more susceptible to damage in high light than wild type plants which immediately proves the central importance of this mechanism under changing environmental conditions (Kasahara *et al.*, 2002). However, under ambient light, i.e. 100 μmol photons m^−2^ s^−1^, and long day growth conditions neither we (data not shown) nor others observed any effect on biomass accumulation (Kasahara *et al.*, 2002). In contrast, for plants grown under short day conditions the accumulation response, which is also defective in *chup1* mutants, was shown to enhance leaf photosynthesis and plant biomass production (Gotoh *et al.*, 2018). In line with these observations, we found similar rates of net CO_2_ assimilation in Col-0 and *chup1* at growth PAR intensity under ambient and low temperature (see Fig. 2 A) while Gotoh and colleagues observed a significantly reduced rate of net CO_2_ assimilation in *chup1* grown under short day. Together with the finding of lowered net CO_2_ assimilation rates in *chup1* under high light intensity, i.e. 1200 μmol photons m^−2^ s^−1^, these results indicate that *chup1* plants can compensate for effects on photosynthetic CO_2_ uptake rates under long photoperiods.

Under control long day growth conditions, neither of the recorded chlorophyll fluorescence parameters significantly differed between Col-0 and *chup1* which was also observed previously under short day growth conditions (Gotoh *et al.*, 2018). This suggests that potential limitation of CO_2_ net assimilation rates in *chup1* rather arises from biochemistry than photochemistry. Supportively, *chup1* assimilated significantly less CO_2_ than Col-0 under ambient temperature, high light intensity and lowered CO_2_ concentration (200ppm, Figure 2 B and Supplementary Table II). While under very low CO_2_ concentrations, i.e. 50ppm, no significant effect was observed this might be due to the very strong overlaying effect of such a condition on all photosynthetic parameters, also in Col-0. Further evidence for a biochemical limitation of photosynthesis in *chup1* is provided by the finding that dynamics of enzyme activities, particularly of SPS, GlcK and Frck, were significantly lowered in *chup1* compared to Col-0 when exposed to 4°C (see Fig. 5). Previous studies have shown a central role of sucrose biosynthesis for cold acclimation and stabilization of photosynthesis (Strand *et al.*, 2003, Nägele *et al.*, 2012). Yet, although the median SPS activity in Col-0 was ~1.5-fold higher than in *chup1* under AL/LT, neither simulated steady state SPS flux nor Suc amount differed between both genotypes (Figs. 3 and 6). This might be explained by the significantly higher amount of F6P in *chup1* than in Col-0 under AL/LT (see Fig. 4). Together with UDP glucose, F6P is substrate for the SPS reaction and, thus, our data suggest that a higher amount of F6P may compensate for a reduced SPS activity to results in a similar rate of Suc biosynthesis (rSPS) in both genotypes. However, our modelling approach comprised simplifications which need to be discussed and addressed in further studies in order to reveal how far estimated reaction rates reflect actual *in vivo* rates. First, in the present study we did not quantify UDP glucose but instead assumed its amount to be ~20% of G6P as reported before (Szecowka *et al.*, 2013). Further, we ignored subcellular distribution of metabolites and, thus, simulated reaction rates might overestimate actual reaction rates due to a higher substrate concentration. For example, approximately one third of F6P amount has previously been reported to be located in the chloroplast of Col-0 and two thirds are located in the cytosol (Szecowka *et al.*, 2013). Future studies on subcellular metabolite distribution might refine the estimated reaction rates and reveal whether compartmentation differs between Col-0 and *chup1*.

While rSPS was similar in both genotypes under all conditions, rates of sucrose cleavage (rINV), fructose phosphorylation (rFRCK) and glucose phosphorylation (rGLCK) were lower in *chup1* than in Col-0 under control and AL/LT conditions (see Fig. 6). Together with rSPS, these reactions describe a cyclic reaction of Suc synthesis and degradation which has been termed as a *futile cycle* due to its seemingly futile consumption of ATP (Geigenberger and Stitt, 1991, Nguyen-Quoc and Foyer, 2001). However, it has also been discussed earlier that the “waste” of ATP might only comprise 3-5% while, as a benefit, such futile cycles increase sensitivity of metabolic regulation (Hue, 1981, Geigenberger and Stitt, 1991). Also, in our own studies we found evidence for a central role of sucrose cycling in stabilization of photosynthesis during cold exposure (Nägele and Heyer, 2013, Weiszmann *et al.*, 2018). Interestingly, while most of the recorded photosynthetic parameters under AL/LT were similar in both genotypes, maximum quantum yield was significantly affected, i.e. lower, in *chup1* which correlated with rINV, rGLCK and rFRCK. Thus, while *chup1* did not suffer from severe photodamage under AL/LT, our findings provide further evidence for an important role of sucrose cycling in tight regulation of photosynthesis under low temperature. Particularly for the study of metabolic cycles mathematical modelling is the method of choice due to underlying complexity, nested uncertainties and non-intuitive modes of regulation (Schaber *et al.*, 2009, Reznik and Segrè, 2010, Henkel *et al.*, 2011).

In addition to activity of hexokinase, oxidation of G6P, catalysed by G6PDH, represents a further important mechanism involved in dynamics of hexose phosphate concentrations under changing environmental conditions. Cold acclimation under AL/LT resulted in a ~3-fold increase of the G6P/F6P ratio in Col-0 and only ~1.5-fold in *chup1*. *Vice versa*, under LL/LT this ratio increased ~3-fold in *chup1* compared to control while in Col-0 concentrations of G6P and F6P increased similarly. A significant increase of the G6P/F6P ratio due to cold acclimation has been reported earlier (Savitch *et al.*, 2001). As suggested by Savitch and colleagues, induction of the ascorbate-glutathione cycle as a ROS defence system might results in an increased supply of NADPH + H^+^ by an increased activity of the oxidative pentose phosphate pathway, oxPPP (May *et al.*, 1998, Savitch *et al.*, 2001). In *chup1*, both G6P and F6P amount significantly increased under AL/LT which together with the wild type-like activities of G6PDH and no obvious photodamage might indicate that ROS defence via oxPPP is still functional. Yet, overaccumulation of G6P under LL/LT and higher rG6PDH rates indicate a less tightly regulated balance between photosynthetic light reactions and carbohydrate metabolism in *chup1* which, under high light intensities, would result in lowered CO_2_ assimilation rates and photodamage.

## Conclusion

Although cold acclimation of *chup1* did not result in a lowered photosynthetic CO_2_ assimilation rate compared to Col-0, the presented findings suggest a central role of CHUP1-mediated chloroplast positioning for tight regulation of photosynthesis and carbohydrate metabolism under low temperature. Future studies might address metabolic effects on the subcellular level, including further mutants affected in chloroplast positioning, e.g. *phototropin* mutants. While the combination of high light intensities and low temperature leads to photoinhibition which prevents the study of metabolic regulation under such conditions, further insights into the effects of chloroplast positioning on metabolic regulation might be gained using other abiotic factors, e.g. heat, high light and a combination of both.

## Acknowledgements

We thank the groups of Plant Evolutionary Cell Biology and Plant Development at LMU München for many fruitful discussions and support.

## Author contributions

AK performed experiments, KS performed microscopy, LF, LS, TS, SF and TN supported experimental analysis. AK and TN analysed data and wrote the manuscript. All authors approved the manuscript.

## Supplementary Information

**Supplementary Figure S1.**
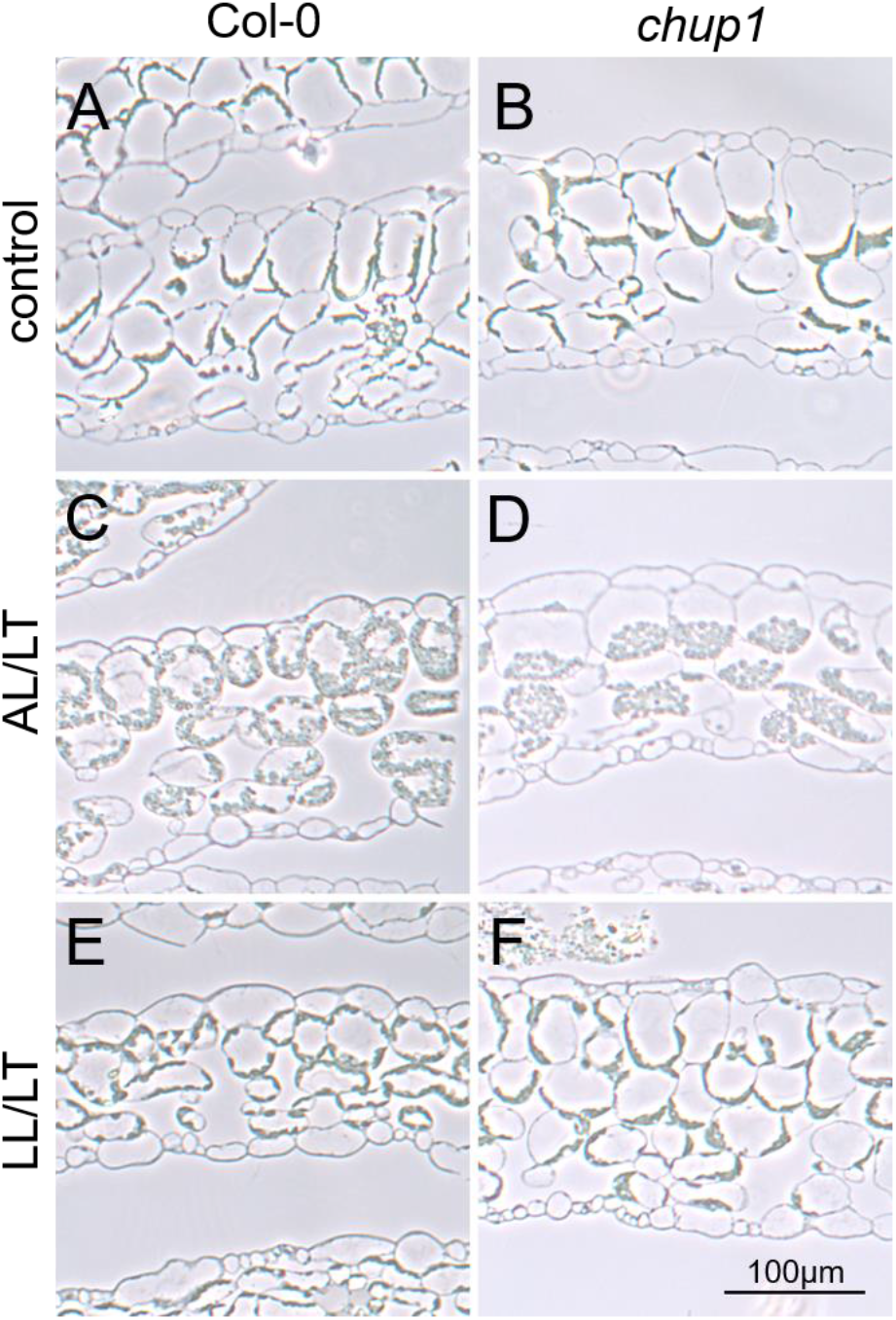
Semi-thin sections of Col-0 and *chup1* primary leaves. (**A, C, E**) Col-0 under control, AL/LT and LL/LT conditions. (**B, D, F**) *chup1* under control, AL/LT and LL/LT conditions. At the end of the night, leaves were kept further 6 hours in darkness to reduce starch content before fixation and ultra-thin-sectioning. Scale bar: 100μm.

**Supplementary Figure S2.**
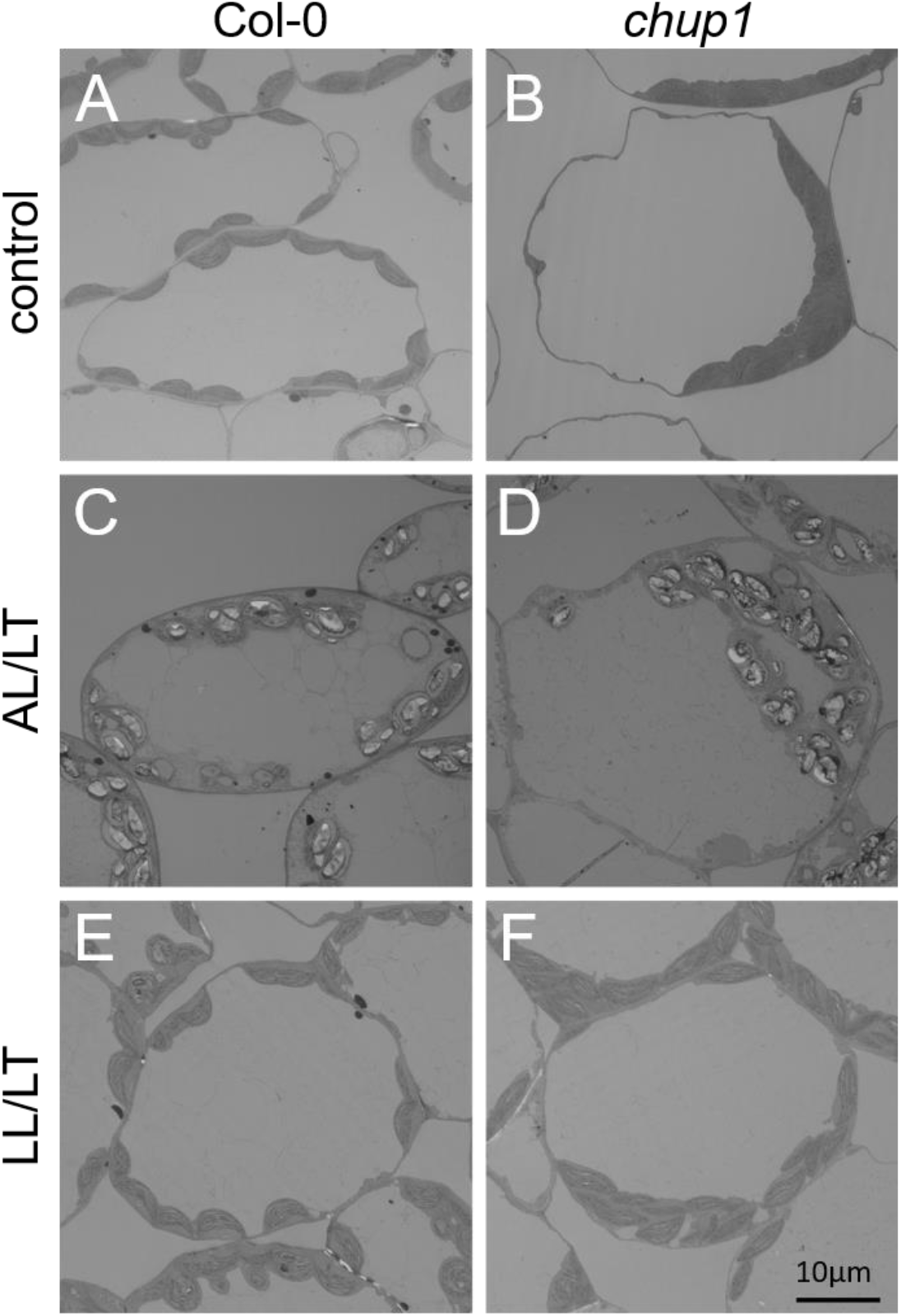
Ultrastructure of mesophyll leaf cells in Col-0 and *chup1* primary leaves. (**A, C, E**) Col-0 under control, AL/LT and LL/LT conditions. (**B, D, F**) *chup1* under control, AL/LT and LL/LT conditions. At the end of the night, leaves were kept further 6 hours in darkness to reduce starch content before fixation and ultra-thin-sectioning. Scale bar: 10μm.

**Supplementary Figure S3.**
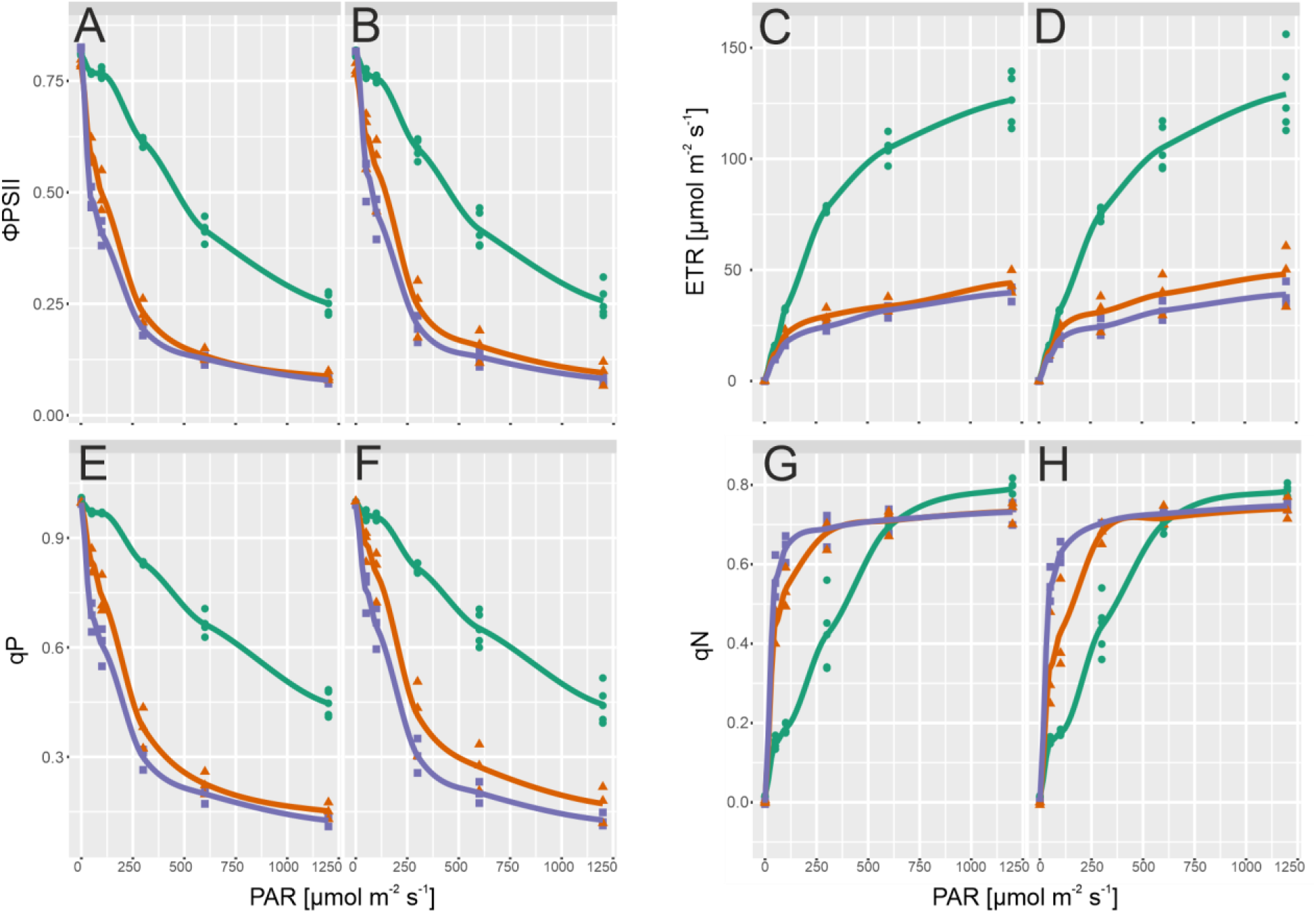
Light response curves of chlorophyll fluorescence parameters. Effective quantum yield of (A) Col-0 and (B) *chup1*. Electron transport rate of (C) Col-0 and (D) *chup1*. Photochemical quenching of (E) Col-0 and (F) *chup1*. Non-photochemical quenching of (G) Col-0 and (H) *chup1*. Green lines and circles: control. Orange lines and triangles: AL/LT. Purple lines and squares: LL/LT. Symbols represent measurements of independent biological replicates (n≥3). Lines represent a local polynomial regression. Measurements of control samples were performed at 22°C, measurements of AL/LT and LL/LT at 4°C.

**Supplementary Figure S4.**
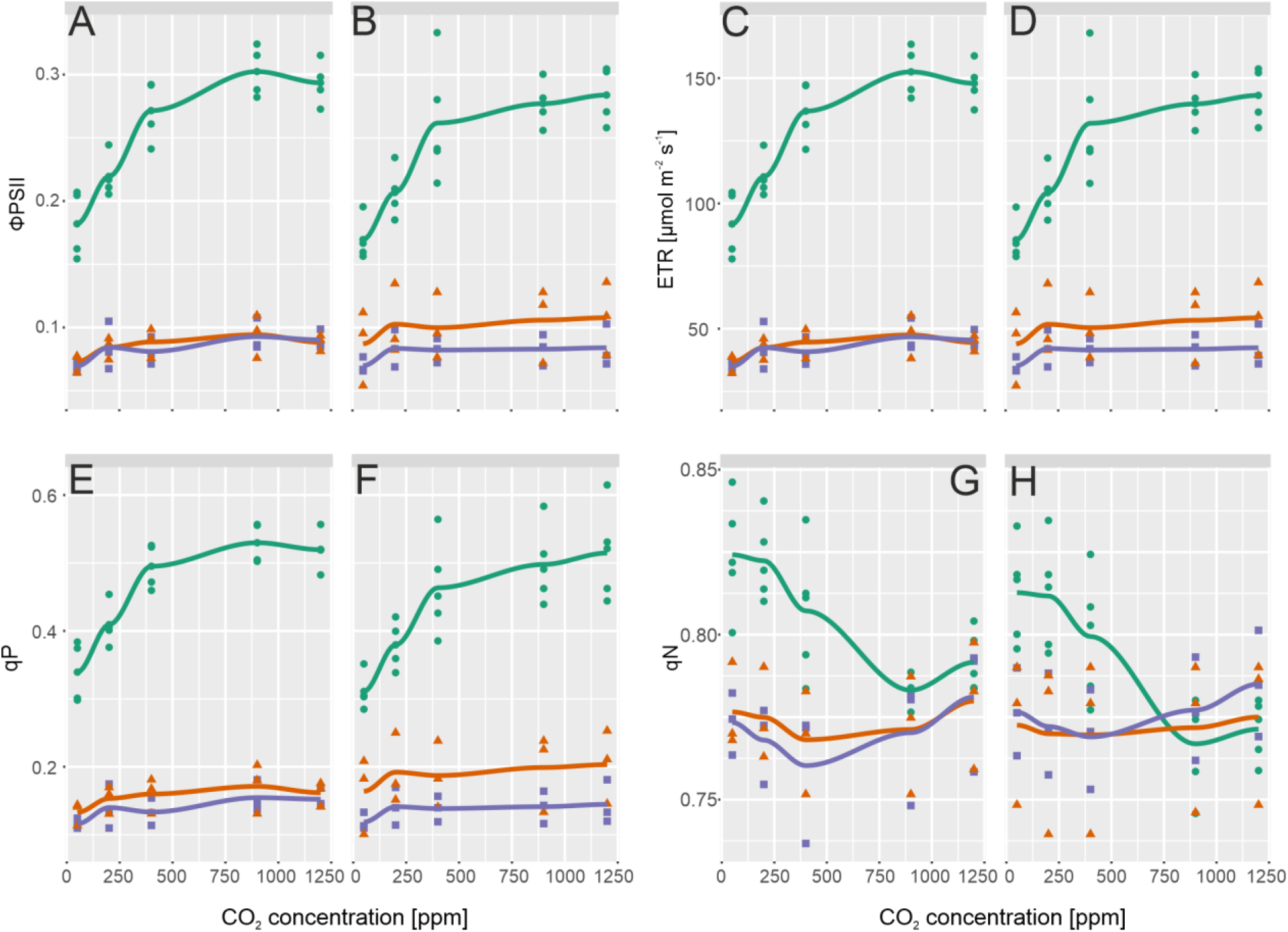
CO_2_ response curves of chlorophyll fluorescence parameters. Effective quantum yield of (**A**) Col-0 and (**B**) *chup1*. Electron transport rate of (**C**) Col-0 and (**D**) *chup1*. Photochemical quenching of (**E**) Col-0 and (**F**) *chup1*. Non-photochemical quenching of (**G**) Col-0 and (**H**) *chup1*. Green lines and circles: control. Orange lines and triangles: AL/LT. Purple lines and squares: LL/LT. Symbols represent measurements of independent biological replicates (n≥3). Lines represent a local polynomial regression. Measurements of control samples were performed at 22°C, measurements of AL/LT and LL/LT at 4°C.

**Supplementary Figure S5.**
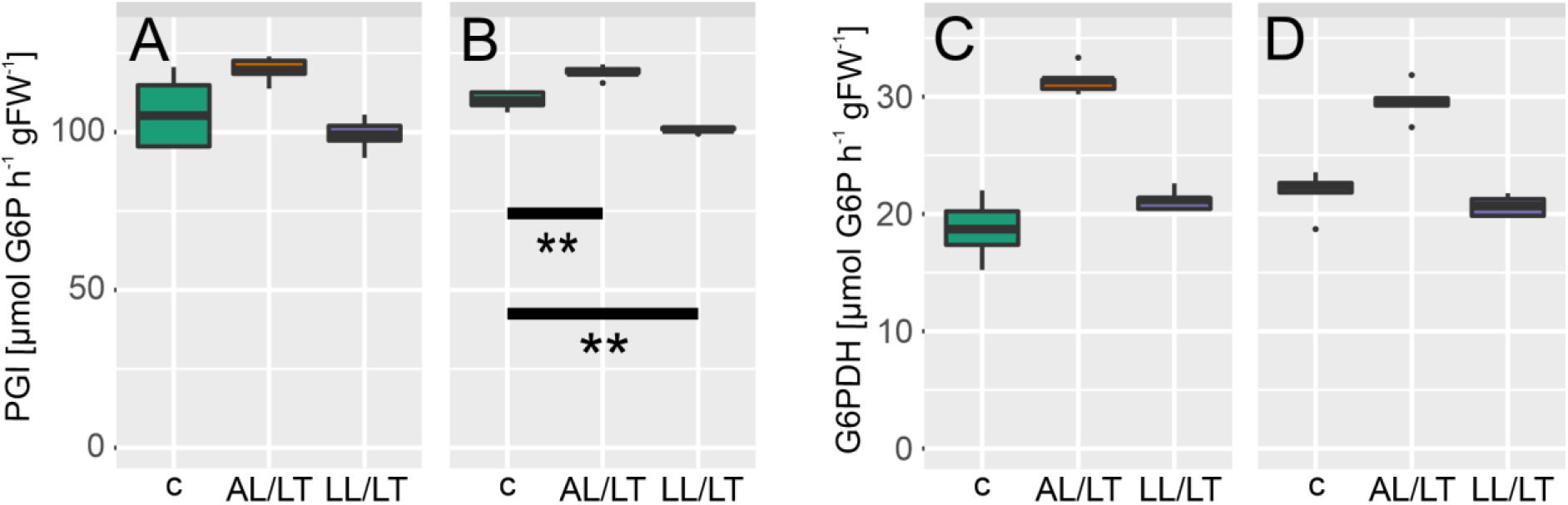
Cold-induced dynamics of PGI and G6PDH activity. **(A, C)** enzyme activities in Col-0, **(B, D)** enzyme activities in *chup1*. Boxes in each panel, left/green: control; middle/orange: AL/LT; right/purple: LL/LT. PGI: phosphoglucoisomerase; G6PDH: glucose 6-phosphate dehydrogenase. Significances are indicated by asterisks but only shown where different between Col-0 and *chup1* (ANOVA; ** p <0.01; *** p<0.001). A complete overview of significances is provided in the supplements (Supplementary Table I). n = 5.

**Supplementary Figure S6.**
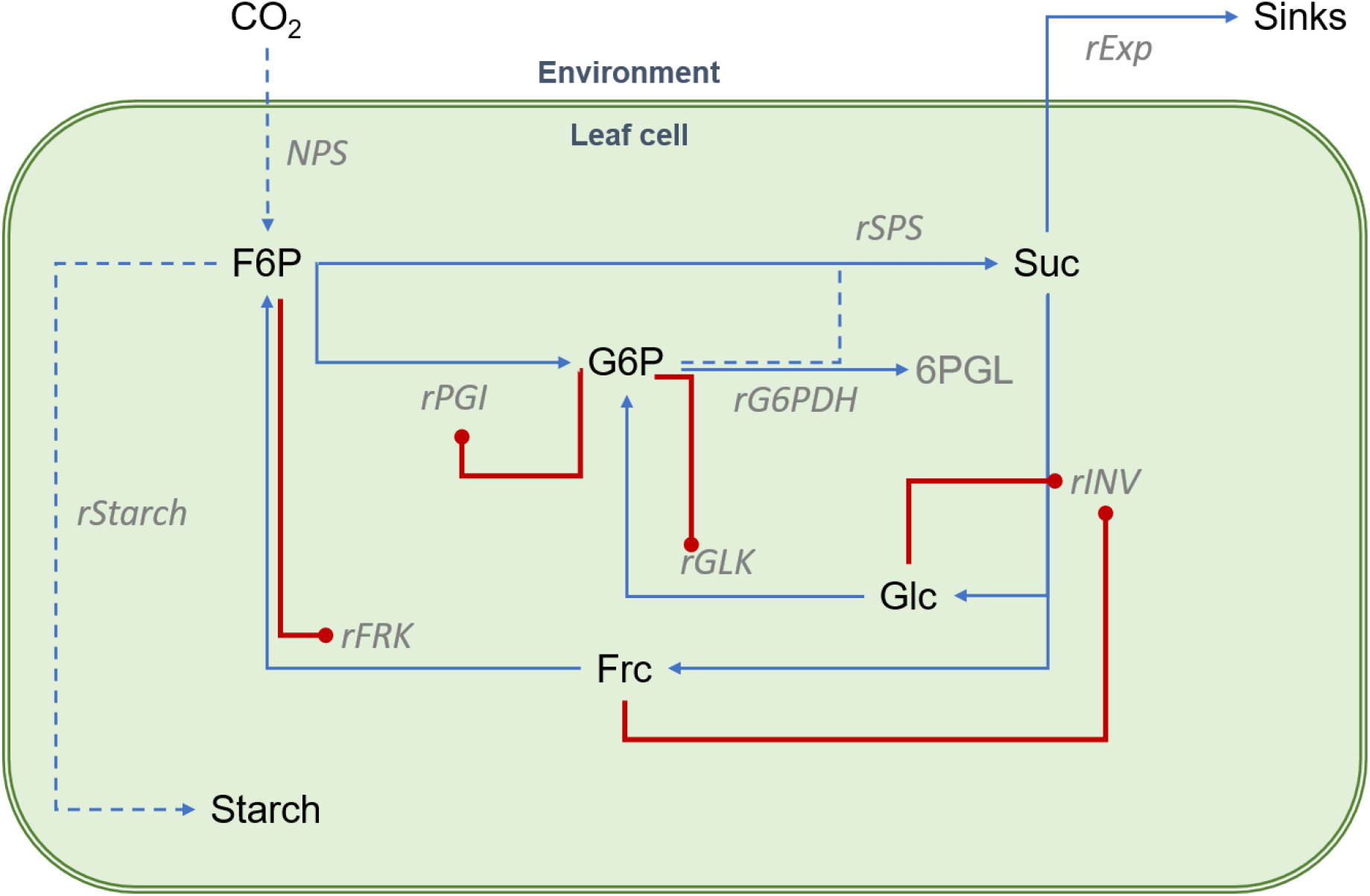
Schematic structure of the central carbohydrate metabolism. Arrows represent enzyme reactions which were simulated by solving a kinetic model based on ordinary differential equations. Red lines indicate inhibition. Dotted lines indicate lumped reactions. Further information about ODEs, reaction kinetics and parameters is provided in the supplements. F6P: fructose 6-phosphate; G6P: glucose 6-phosphate; 6PGL: 6-phosphogluconolactone; Suc: sucrose; Glc: glucose; Frc: fructose; NPS: net photosynthesis; rStarch: rate of net starch synthesis. rSPS: reaction rate of sucrose phosphate synthase; rINV: reaction rate of invertase; rGLK: reaction rate of glucokinase; rFRK: reaction rate of fructokinase; rPGI: reaction rate of phosphoglucose isomerase; rG6PDH: reaction rate of glucose 6-phosphate dehydrogenase; rEXP: rate of sucrose export to sinks.

**Supplementary Table I.**
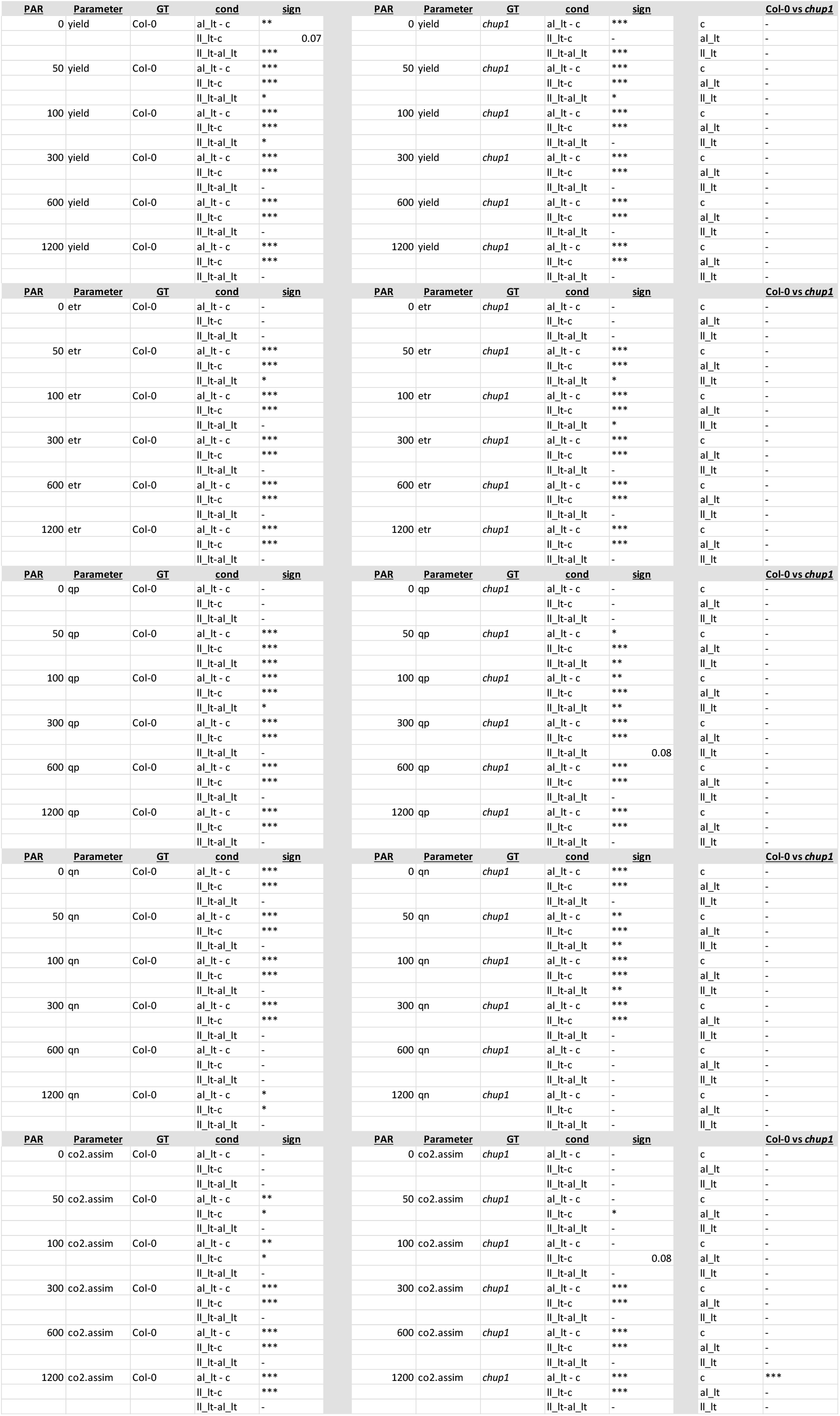
ANOVA results of photosynthesis light response curves. c: control; al_lt: ambient light low temperature; ll_lt: low light low temperature. * p<0.05; ** p<0.01; *** p<0.001.

**Supplementary Table II.**
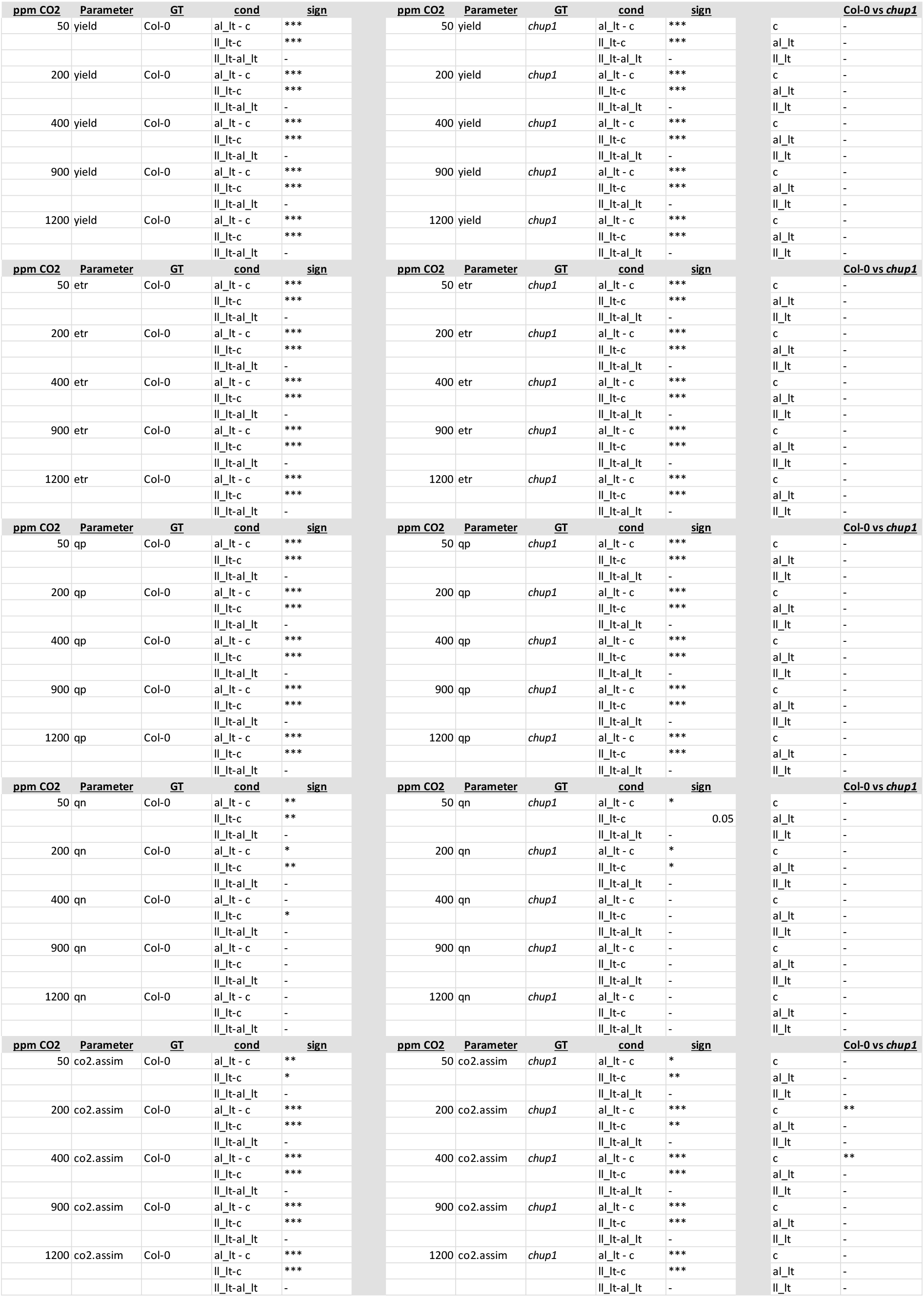
ANOVA results of photosynthesis CO_2_ response curves. c: control; al_lt: ambient light low temperature; ll_lt: low light low temperature. * p<0.05; ** p<0.01; *** p<0.001.

**Supplementary Table III.**
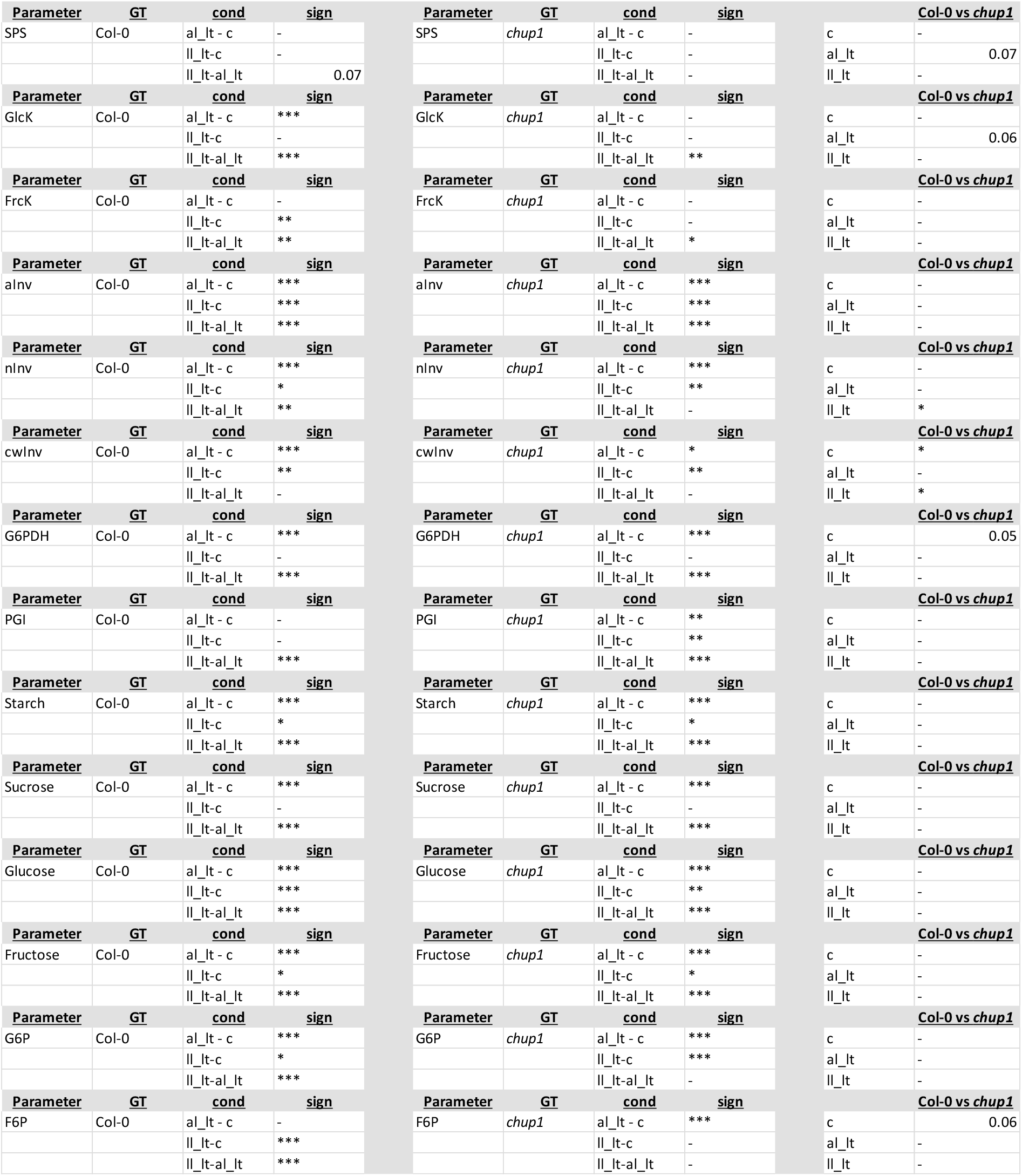
ANOVA results of molecular data. c: control; al_lt: ambient light low temperature; ll_lt: low light low temperature. * p<0.05; ** p<0.01; *** p<0.001.

**Supplementary Table IV:** Ordinary differential equations and enzyme kinetics.

|  |  |  |  |  |  |
| --- | --- | --- | --- | --- | --- |
| **<u>ODE system of the kinetic model</u>** |  |  |  |  |  |
| $d/dt(F6P) = (1/6)*rNPS - rStarch - rSPS + rFRCK - rPGI$ | | | | | |
| $d/dt(G6P) = rPGI - rSPS + rGLCK - rG6PDH$ | | | | | |
| $d/dt(Suc) = rSPS - rINV - rEXP$ | | | | | |
| $d/dt(Glc) = rINV - rGLCK$ | | | | | |
| $d/dt(Frc) = rINV - rFRCK$ | | | | | |
| **<u>Model reactions and kinetic equations</u>** |  |  |  |  |  |
| $rPGI = (vmax\_pgi*F6P)/(Km\_pgi*(1+G6P/Ki\_pgi\_g6p)+F6P)$ | | | | | |
| $rG6PDH = (vmax\_g6pdh*G6P)/(Km\_g6pdh+G6P)$ | | | | | |
| $rSPS = vmax\_sps*(F6P/(Km\_sps\_f6p+F6P))*(0.2*G6P/(Km\_sps\_g6p+0.2*G6P))$ | | | | | |
| $rGLCK = (vmax\_glck*Glc)/(Km\_glck*(1+G6P/Ki\_glck\_g6p) + Glc)$ | | | | | |
| $rFRCK = (vmax\_frck*Frc)/(Km\_frck*(1+F6P/Ki\_frck\_f6p) + Frc)$ | | | | | |
| $rINV = (vmax\_inv*Suc)/((Km\_inv*(1+Frc/Ki\_inv\_frc)+Suc)*(1+Glc/Ki\_inv\_glc))$ | | | | | |
| $rEXP = kEXP*Suc$ | | | | | |

**Supplementary Table V:**
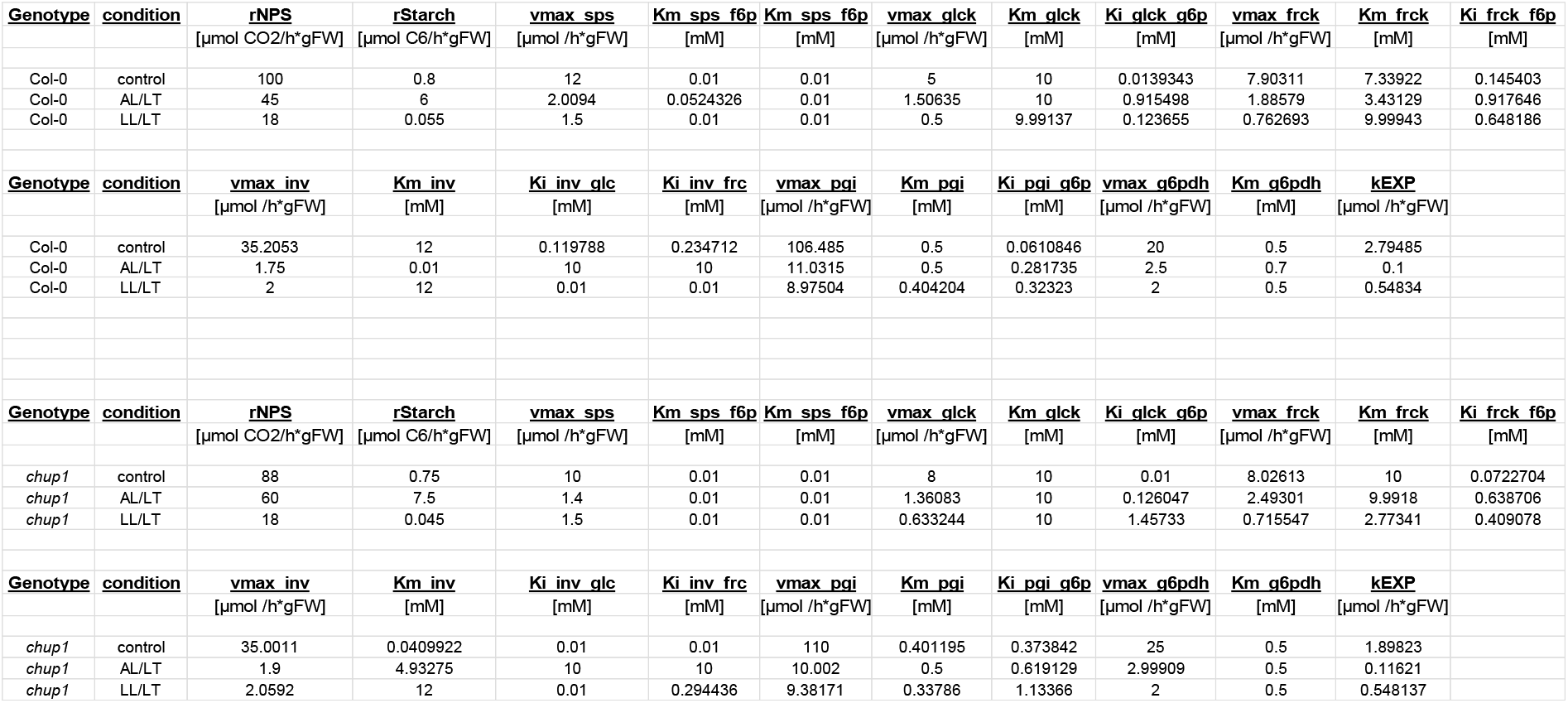
Optimized kinetic parameters for steady state simulations in Col-0 and *chup1*.

**Supplementary Table VI:** Range boundaries of model parameters for parameter optimization. Optimization algorithm: (Global) Particle swarm pattern search (Vaz and Vicente, 2007).

| <u>Col-0 control</u> |  |  | <u>Col-0 AL/LT</u> |  |  | <u>Col-0 LL/LT</u> |  |  |
| --- | --- | --- | --- | --- | --- | --- | --- | --- |
| Parameter | Lower bound | Upper bound | Parameter | Lower bound | Upper bound | Parameter | Lower bound | Upper bound |
| rStarch | 0.7 | 0.8 | rStarch | 6 | 6.5 | rStarch | 0.045 | 0.055 |
| vmax_sps | 8 | 12 | vmax_sps | 2 | 2.5 | vmax_sps | 1 | 1.5 |
| Km_sps_f6p | 0.01 | 10 | Km_sps_f6p | 0.01 | 10 | Km_sps_f6p | 0.01 | 10 |
| Km_sps_f6p | 0.01 | 10 | Km_sps_f6p | 0.01 | 10 | Km_sps_f6p | 0.01 | 10 |
| vmax_glck | 5 | 7 | vmax_glck | 1.5 | 2 | vmax_glck | 0.5 | 0.75 |
| Km_glck | 0.01 | 10 | Km_glck | 0.01 | 10 | Km_glck | 0.01 | 10 |
| Ki_glck_g6p | 0.01 | 10 | Ki_glck_g6p | 0.01 | 10 | Ki_glck_g6p | 0.01 | 10 |
| vmax_frck | 7 | 9 | vmax_frck | 1.5 | 2 | vmax_frck | 0.75 | 0.9 |
| Km_frck | 0.01 | 10 | Km_frck | 0.01 | 10 | Km_frck | 0.01 | 10 |
| Ki_frck_f6p | 0.01 | 10 | Ki_frck_f6p | 0.01 | 10 | Ki_frck_f6p | 0.01 | 10 |
| vmax_inv | 35 | 45 | vmax_inv | 1.25 | 1.75 | vmax_inv | 2 | 3 |
| Km_inv | 0.01 | 12 | Km_inv | 0.01 | 12 | Km_inv | 0.01 | 12 |
| Ki_inv_glc | 0.01 | 10 | Ki_inv_glc | 0.01 | 10 | Ki_inv_glc | 0.01 | 10 |
| Ki_inv_frc | 0.01 | 10 | Ki_inv_frc | 0.01 | 10 | Ki_inv_frc | 0.01 | 10 |
| vmax_pgi | 100 | 110 | vmax_pgi | 10 | 12 | vmax_pgi | 8.5 | 9.5 |
| Km_pgi | 0.3 | 0.5 | Km_pgi | 0.3 | 0.5 | Km_pgi | 0.3 | 0.5 |
| Ki_pgi_g6p | 0.0001 | 10 | Ki_pgi_g6p | 0.0001 | 10 | Ki_pgi_g6p | 0.0001 | 10 |
| vmax_g6pdh | 17 | 20 | vmax_g6pdh | 2.5 | 3.5 | vmax_g6pdh | 1.9 | 2 |
| Km_g6pdh | 0.5 | 0.7 | Km_g6pdh | 0.5 | 0.7 | Km_g6pdh | 0.5 | 0.7 |
| kEXP | 0.1 | 10 | kEXP | 0.1 | 10 | kEXP | 0.1 | 10 |
| <u>chup1 control</u> |  |  | <u>chup1 AL/LT</u> |  |  | <u>chup1 LL/LT</u> |  |  |
| Parameter | Lower bound | Upper bound | Parameter | Lower bound | Upper bound | Parameter | Lower bound | Upper bound |
| rStarch | 0.66 | 0.75 | rStarch | 6.5 | 7.5 | rStarch | 0.035 | 0.045 |
| vmax_sps | 9 | 10 | vmax_sps | 1.2 | 1.4 | vmax_sps | 1 | 1.5 |
| Km_sps_f6p | 0.01 | 10 | Km_sps_f6p | 0.01 | 10 | Km_sps_f6p | 0.01 | 10 |
| Km_sps_f6p | 0.01 | 10 | Km_sps_f6p | 0.01 | 10 | Km_sps_f6p | 0.01 | 10 |
| vmax_glck | 8 | 9 | vmax_glck | 1.3 | 1.5 | vmax_glck | 0.6 | 1 |
| Km_glck | 0.01 | 10 | Km_glck | 0.01 | 10 | Km_glck | 0.01 | 10 |
| Ki_glck_g6p | 0.01 | 10 | Ki_glck_g6p | 0.01 | 10 | Ki_glck_g6p | 0.01 | 10 |
| vmax_frck | 8 | 9 | vmax_frck | 1.5 | 2.5 | vmax_frck | 0.7 | 1.1 |
| Km_frck | 0.01 | 10 | Km_frck | 0.01 | 10 | Km_frck | 0.01 | 10 |
| Ki_frck_f6p | 0.01 | 10 | Ki_frck_f6p | 0.01 | 10 | Ki_frck_f6p | 0.01 | 10 |
| vmax_inv | 35 | 45 | vmax_inv | 1.7 | 1.9 | vmax_inv | 2 | 3 |
| Km_inv | 0.01 | 12 | Km_inv | 0.01 | 12 | Km_inv | 0.01 | 12 |
| Ki_inv_glc | 0.01 | 10 | Ki_inv_glc | 0.01 | 10 | Ki_inv_glc | 0.01 | 10 |
| Ki_inv_frc | 0.01 | 10 | Ki_inv_frc | 0.01 | 10 | Ki_inv_frc | 0.01 | 10 |
| vmax_pgi | 105 | 115 | vmax_pgi | 10 | 12 | vmax_pgi | 9 | 9.5 |
| Km_pgi | 0.3 | 0.5 | Km_pgi | 0.3 | 0.5 | Km_pgi | 0.3 | 0.5 |
| Ki_pgi_g6p | 0.0001 | 10 | Ki_pgi_g6p | 0.0001 | 10 | Ki_pgi_g6p | 0.0001 | 10 |
| vmax_g6pdh | 20 | 25 | vmax_g6pdh | 2.5 | 3 | vmax_g6pdh | 1.8 | 2 |
| Km_g6pdh | 0.5 | 0.7 | Km_g6pdh | 0.5 | 0.7 | Km_g6pdh | 0.5 | 0.7 |
| kEXP | 0.1 | 10 | kEXP | 0.1 | 10 | kEXP | 0.1 | 10 |

